# Leveraging high-powered RNA-Seq datasets to improve inference of regulatory activity in single-cell RNA-Seq data

**DOI:** 10.1101/553040

**Authors:** Ning Wang, Andrew E. Teschendorff

## Abstract

Inferring the activity of transcription factors in single cells is a key task to improve our understanding of development and complex genetic diseases. This task is, however, challenging due to the relatively large dropout rate and noisy nature of single-cell RNA-Seq data. Here we present a novel statistical inference framework called SCIRA (Single Cell Inference of Regulatory Activity), which leverages the power of large-scale bulk RNA-Seq datasets to infer high-quality tissue-specific regulatory networks, from which regulatory activity estimates in single cells can be subsequently obtained. We show that SCIRA can correctly infer regulatory activity of transcription factors affected by high technical dropouts. In particular, SCIRA can improve sensitivity by as much as 70% compared to differential expression analysis and current state-of-the-art methods. Importantly, SCIRA can reveal novel regulators of cell-fate in tissue-development, even for cell-types that only make up 5% of the tissue, and can identify key novel tumor suppressor genes in cancer at single cell resolution. In summary, SCIRA will be an invaluable tool for single-cell studies aiming to accurately map activity patterns of key transcription factors during development, and how these are altered in disease.

## Introduction

Transcription factors play key roles in cellular development, and the regulatory networks they form are often disrupted in complex genetic disease. Elucidating the regulatory networks orchestrating cellular development is a challenging endeavor, in part because inferring regulatory relations and activity has so far been done mostly in the context of bulk cell population level data where regulatory influences may be averaged out ^1,2^. Regulatory relationships function at the single-cell level and therefore their implications can only be truly understood at single-cell resolution ^3–8^.

Recent single-cell RNA-sequencing (scRNA-Seq) studies have significantly advanced our understanding of cellular development and disease ^4,7, 9–29^, yet how best to infer regulatory activity at the single-cell level is still unclear. While a number of methods for inferring regulatory activity have appeared ^30–32^, all attempt to do so by first inferring the regulatory interactions from the noisy scRNA-Seq data itself, an approach recently assessed by others to be highly suboptimal ^33^, mainly because of the relatively high dropout rate of such data ^5, 34^. The high dropout rate of scRNA-Seq data means that for low to moderately expressed TFs, inferring their differential activity may not be possible from its measured mRNA level alone, thus precluding direct application of methods that attempt to reverse engineer regulatory relationships ^35^. As a result, currently available methods will in general lack the sensitivity to identify transcription factors (TFs) that play a key role in development, or to identify which specific TFs are disrupted in disease. Thus, novel algorithms are needed that can infer regulatory activity in the context of high dropout rate event data, and that can do so at substantially improved sensitivity over current state-of-the-art methods.

Here we present a novel algorithm called SCIRA (Single Cell Inference of Regulatory Activity), which improves the sensitivity to detect true transcription factor activity patterns by as much as 70% or more, over the current state-of-the-art. Our work supports the view that regulatory relationships can’t in general be reliably inferred from scRNA-Seq data, pointing towards the need for a hybrid approach which leverages the power of large-scale high quality bulk RNA-Seq datasets to infer an approximate regulatory influence network, which can subsequently be used to infer regulatory activity in single-cells. SCIRA is such a hybrid approach which uses the inferred predicted targets (or “regulon”) of a given TF from bulk data to infer its regulatory activity in each single cell. Importantly, we demonstrate that the use of an appropriately powered bulk tissue dataset to infer regulons does not present a limitation, as SCIRA is able to identify novel regulators of cell-fate in single-cell data, even for cell-types that only make up 5% of cells in a bulk tissue sample.

## Results

### Motivation and rationale for the SCIRA algorithm

The inference of regulatory activity in a single cell is a task of great importance. Because scRNA-Seq is noisy and subject to a high technical dropout rate ^5^, and given a recent report which concluded that inferring regulatory relationships from scRNA-Seq is a highly suboptimal procedure ^33^, we decided to approach this task differently, by first using a large-scale bulk RNA-Seq dataset to construct an approximate regulatory network. In particular, we here use the GTEX dataset ^36^, which consists of bulk RNA-Seq profiles for 8555 samples encompassing 30 tissue types (**Methods**), to infer a regulatory network with our SEPIRA algorithm ^37^ (**Methods, Fig.1A**). SEPIRA uses a greedy partial correlation framework, similar in spirit to the state-of-the-art GENIE3 algorithm ^35, 38^, to infer direct dependencies between regulators and downstream targets. SEPIRA constructs these regulons for tissue-specific TFs only, whilst also adjusting for cell-type (stromal) heterogeneity, which can otherwise strongly confound correlation and differential expression analyses ^39^. We have estimated, using power-analysis, that SCIRA has approximately 60-70% sensitivity to capture tissue-specific TFs that are however only highly expressed in 5% of the cells in the tissue (**SI fig.S1**, **Methods**). Correspondingly, sensitivities are close to 100% for TFs highly expressed in a cell-type that makes up at least 20% of the cells in the tissue (**SI fig.S1**). Thus, by leveraging the high-power of the GTEX dataset, SEPIRA builds a high-confidence tissue-specific regulatory network, including TFs that mark under-represented cell-types in the tissue (**Fig.1A**), with a view to subsequently apply the regulons of all derived TFs to scRNA-Seq datasets profiling cells in the same tissue-type (**Fig.1B**). By using the actual regulon of the TF as a target profile in a linear regression model framework ^40^ (**Methods**), inference of the regulatory activity of the TF should be robust to dropouts, even if the TF itself is not detected across most if not all of the cells in the study (**Fig.1B**). We show below that a very high technical dropout rate can affect even TFs that are known to be highly expressed in the given tissue, highlighting the general inadequacy of current scRNA-Seq datasets for inferring regulatory relations. Finally, by estimating regulatory activity of the TFs in each cell one can construct TF-activity maps which can subsequently be analysed with say clustering algorithms to reveal novel insights at single-cell resolution (**Fig.1C**). We refer to the whole algorithmic pipeline above as SCIRA (Single Cell Inference of Regulatory Activity).

**Figure-1:**
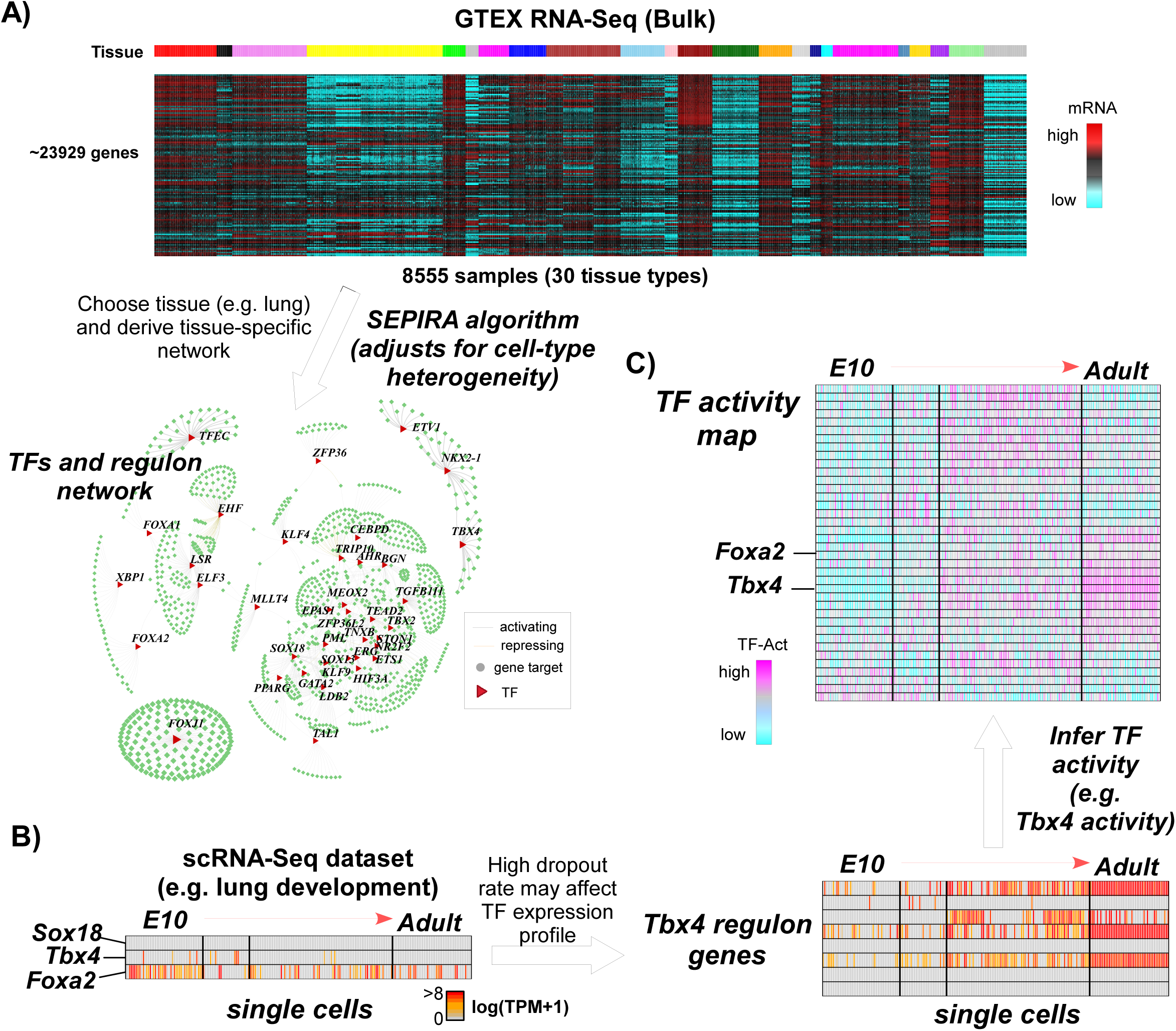
SCIRA workflow. **A)** Since bulk RNA-Seq data does not suffer from technical dropouts and is much more reliable than scRNA-Seq data, for a given choice of tissue, we use the GTEX bulk RNA-Seq expression set to derive a corresponding tissue-specific regulatory network, consisting of tissue-specific transcription factors (TFs) and their targets (regulons). The inference of the network proceeds with the SEPIRA algorithm which used a greedy partial correlation approach, whilst also adjusting for stromal contamination within the tissue. **B)** Given the high technical dropout rate and overall noisy nature of scRNA-Seq data, it is not possible to reliably infer TF activity from the TF expression profile alone. However, using the TF regulons derived in A), and using the genes within the regulon that are not strongly affected by dropouts, we can estimate TF activity across the single-cells. Depicted is an example with 3 lung-specific TFs (Sox18, Tbx4, *Foxa2*) and timepoints representing different developmental stages (e.g. E10=embryonic day 10), as well as the expression pattern of the regulon genes for *Tbx4.* **C)** We use linear regressions between the expression values of all the genes in a given cell and the corresponding TF-regulon profile, to derive the activity of the TF as the t-statistic of the estimated regression coefficient, resulting in a TF activity map over the tissue-specific TFs and single cells.

### Validation and improved sensitivity of SCIRA in liver tissue

To test SCIRA, we first considered the case of a developmental scRNA-Seq study in liver tissue, in which hepatoblasts differentiate into hepatocytes and cholangiocytes, with a total of 447 single cells collected over the course of 7 timepoints during embryonic development ^41^ (**Methods**). Correspondingly, we applied our SEPIRA algorithm to the large GTEX expression compendium dataset which includes liver tissue, resulting in a liver-specific regulatory network of 22 TFs and 788 targets, with an average number of targets per TF of 41 (range 10 to 151) (**SI table S1, Methods**). Among the 22 liver-specific TFs, were well-known factors, such as *Hnf1a, Hnf4a* and *Foxa1/Foxa2*. We validated the regulons of the 22 TFs in two independent datasets: the RNA-Seq multi-tissue set from the ProteinAtlas ^42^ project, and the Affymetrix multi-tissue expression set from Roth et al ^43^ (**SI figs.S2-3**). Since liver-specific TFs ought to exhibit increased activity during a developmental timecourse from hepatoblasts to hepatocytes and cholangiocytes, SCIRA ought to predict this increased TF activity in the scRNA-Seq study above. We applied SCIRA to the scRNA-Seq study, to infer activity estimates for the 22 liver-specific TFs in each of the 447 single cells, which showed that many of the TFs (e.g. *Hnf1a*) exhibited an increased activity pattern during the time course, in line with their known role in liver specification (**Fig.2A, 2C**). This was not observed had we used the expression values of the TFs themselves (**Fig.2B**). Of the 22 TFs, 16 exhibited a significant increase in activity according to SCIRA (Bonferroni adjusted P<0.05, **Fig.2D**), representing a sensitivity of over 70% (**Fig.2E**), whereas differential expression analysis (**Methods**) only predicted 3 to be upregulated (a sensitivity of only 13%, **Fig.2E**). The improved sensitivity of SCIRA over differential expression analysis is statistically significant (Fisher test P-value = 6e-4, **Fig.2E**). Of note, SCIRA-derived TF-activity estimates also anti-correlated with a highly-validated measure of single-cell potency ^44^, as required since our TFs should be more active in the less potent cells (**SI fig.S4**).

**Figure-2:**
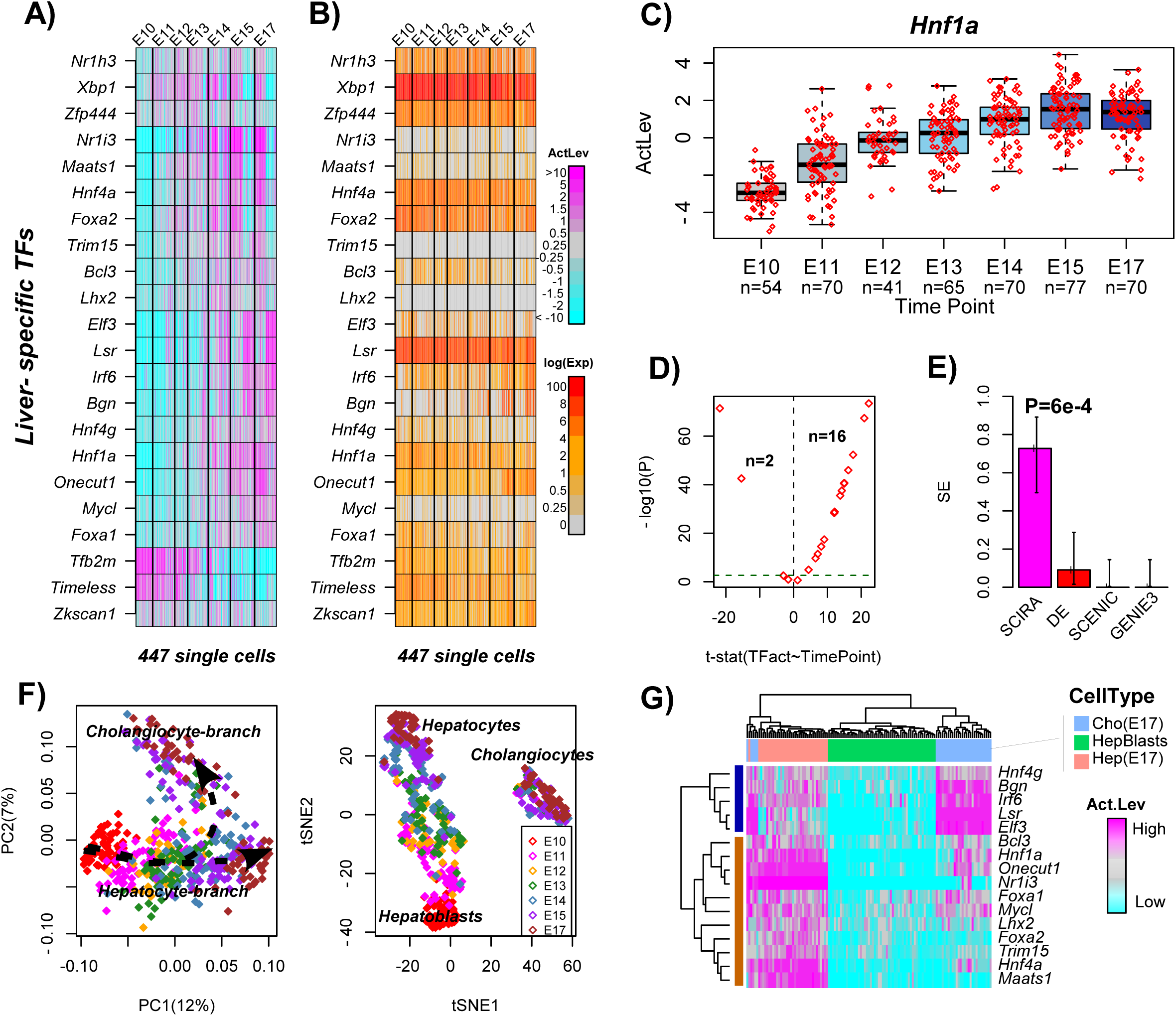
Validation and improved sensitivity of SCIRA in liver development. **A)** Heatmap of regulatory activity of 22 liver-specific TFs across 447 single cells representing seven developmental time points in mouse liver development (e.g E10=embryonic day 10), as estimated using the SCIRA algorithm. **B)** Corresponding heatmap displaying log_2_(TPM+1) values. **C)** Pattern of regulatory activity change for *Hnf1a* as predicted by SCIRA. P-value is from a linear regression. **D)** Scatterplot of t-statistics of a linear regression of activity level against time point (x-axis) vs. the significance of the t-statistic [–log_10_(P-value), y-axis] for all 22 TFs. The number of significant associations at a Bonferroni adjusted P<0.05 level are given. **E)** Barplot comparing the sensitivity (SE) to detect increased activity (SCIRA) or upregulation (DE) across the 22 TFs. Error bars represent 95% confidence intervals estimated using Wilson’s continuity correction. P-value is from a one-tailed Fisher-exact test comparing SCIRA to DE. Also shown are the SE values for SCENIC, and SCENIC without the motif-enrichment analysis step (denoted “GENIE3”). **F)** PCA and t-SNE scatterplots of the 447 single cells obtained by applying PCA and t-SNE on the TF activity matrix shown in A). **G)** Clustering heatmap over the 16 TFs exhibiting increased activity in D), and all single cells at the start (E10) and endpoints (E17). Cell-types annotated as hepatoblasts (E10, Hepblasts), hepatocytes at E17 (Hep(E17)) and cholangiocytes (Cho(E17)).

### SCIRA reveals novel regulators of cholangiocyte fate

Hepatocytes represent the dominant epithelial subtype in liver tissue, with cholangiocytes accounting for only 5% of all liver tissue cells ^45^. We verified that the GTEX liver samples are indeed dominated by hepatocytes and not cholangiocytes (**SI fig.S5**). According to our previous power-analysis, we estimated 60-70% sensitivity to detect cholangiocyte specific TFs, despite cholangiocytes making up such a small proportion of cells in liver tissue. To investigate this, we first performed PCA on the SCIRA-derived transcription factor activity matrix defined over the 22 liver-specific TFs and 447 single cells, which revealed a branching pattern emerging at E14/E15 (**Fig.2F, SI fig.S6**). Using independent markers from MacParland et al ^45^, we can attribute this bifurcation to separate hepatocyte and cholangiocyte branches (**Fig.2F, SI fig.S6**), consistent with previous findings obtained on the full expression matrix (**SI fig.S7**) ^41^. This further validates the regulatory activity estimates obtained with SCIRA. t-stochastic neighborhood embedding (t-SNE) ^46^ analysis helped delineate distinct clusters of single-cells, with cells at the last time point (E17) appearing in both hepatocyte and cholangiocyte branches (**Fig.2F**). Clustering the TF-activity matrix over the 16 TFs which exhibited an increased activity during the time course, and over the single cells collected at the start (E10) and endpoints (E17), revealed three main clusters of cells representing hepatoblasts, differentiated hepatocytes and cholangiocytes (**Fig.2G**). The two clusters of TFs defined those marking hepatocytes (e.g. *Hnf1a, Hnf4a, Foxa1, Foxa2, Onecut1 (Hnf6)*) and cholangiocytes (e.g. *Bgn, Irf6, Lsr, Elf3*). Consistent with this, *Elf3* has recently been shown to be a marker for cholangiocytes ^45^ and *Lsr,* a lipolysis stimulated lipoprotein receptor, is a well-known cholangiocyte marker ^47^, variants of which are associated with pediatric cholestatic liver disease ^48^. Both *Bgn* and *Irf6* also exhibited a bi-modal activation pattern during E14-E15 (**Fig.2A**), the timepoint of branching, suggesting a novel role for these in determining cholangiocyte fate. Thus, SCIRA has been able to detect dynamic patterns of TF activity relevant to cholangiocyte specification, despite their under-representation in the GTEX liver samples.

### Validation and improved sensitivity of SCIRA in lung development

In order to further validate SCIRA, we next considered a well-known single-cell developmental study of the lung epithelium in mice ^17^. A total of 201 cells were profiled using the C1 Fluidigm system and collected from four different developmental timepoints (E14, E16, E18, Adult) (**Methods**). Using SEPIRA on the GTEX data, we constructed a lung-specific regulatory network, termed “LungNet”, which consists of 38 lung-specific TFs and a total of 1145 gene targets (range of targets per TF=10 to 152, mean=39.8) (**SI table.S2**, **Methods**). We note that LungNet was already constructed and extensively validated by us previously ^37^. Since by construction these 38 TFs are more highly expressed in lung tissue compared to all other tissue types and given their established role in lung tissue development ^37^, most if not all of these ought to exhibit increased activation in the scRNA-Seq lung development study. Applying SCIRA, we observed that indeed most of the 38 TFs did exhibit increased activation at some stage during lung development (**Fig.3A, Methods**). One of the top-ranked TFs was *Nkx2-1*, which showed a clear linear increase in activation with developmental stage (**Fig.3C**). Formally, of the 38 TFs, 20 exhibited a significant increase in activation at a Bonferroni corrected P<0.05 level (**Fig.3D**), whilst 26 were associated at a more relaxed unadjusted P<0.05 level. This represents a sensitivity of approximately 50% and 70%, respectively. In line with the increased activity of these TFs with differentiation time point, and therefore with reduced cell potency, most of the TFs exhibited activity patterns that anti-correlated with our signaling entropy potency measure (**SI fig.S7**).

**Figure-3:**
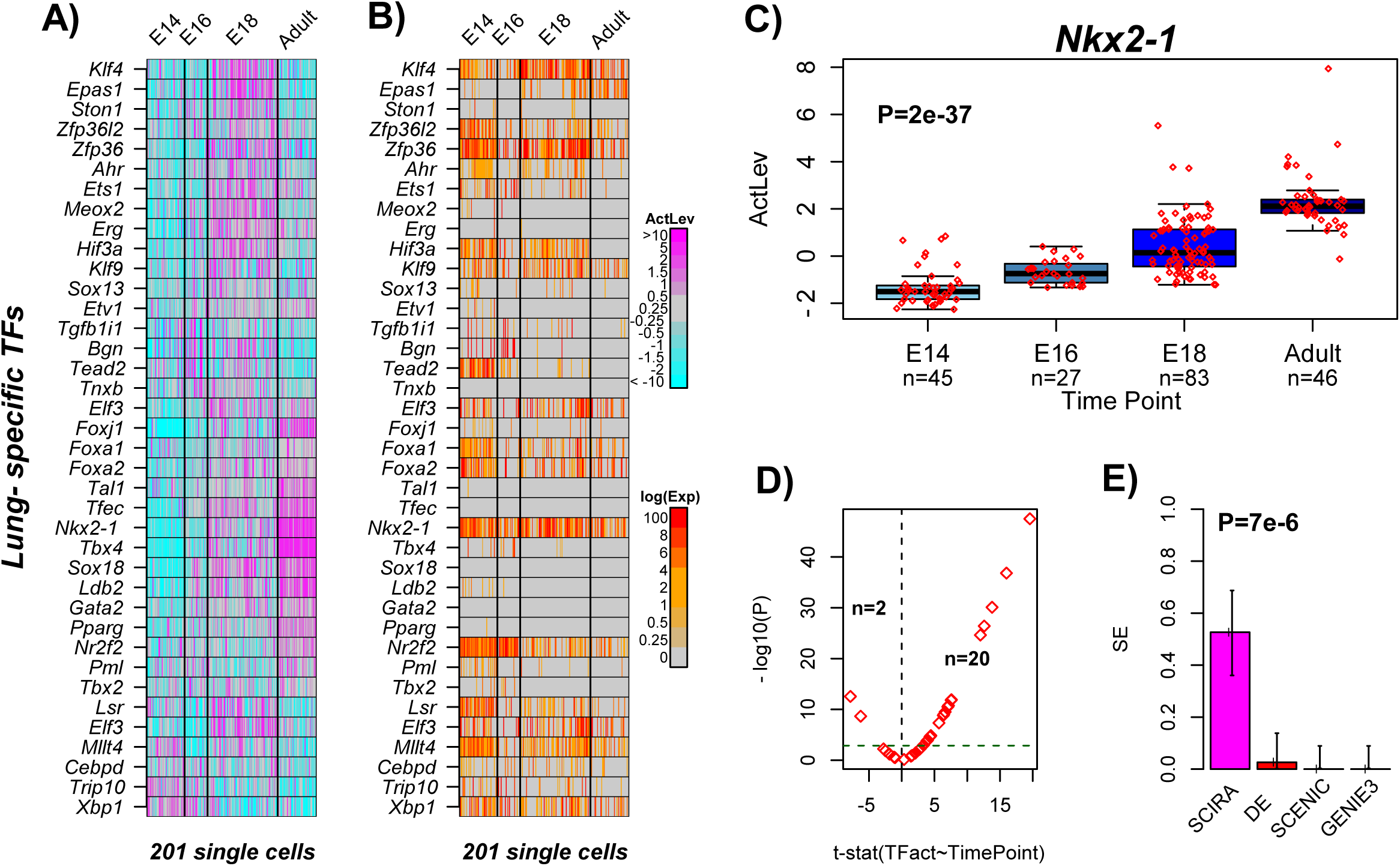
Validation and improved sensitivity of SCIRA in lung development. **A)** Heatmap of regulator activity of 38 lung-specific TFs across 201 single cells representing four developmental time points in lung development in mice, as estimated using the SCIRA algorithm. **B)** Corresponding heatmap displaying log_2_(FPKM+1) values. **C)** Pattern of regulatory activity change for *Nkx2-1.* P-value is from a linear regression. **D)** Scatterplot of t-statistics of a linear regression of activity level against timepoint (x-axis) vs. the significance of the t-statistic [–log_10_(P-value), y-axis] for all 38 TFs. The number of significant associations at a Bonferroni adjusted P<0.05 level are given. **E)** Barplot comparing the sensitivity (SE) to detect increased activity (SCIRA) or upregulation (DE) across the 38 TFs. Error bars represent 95% confidence intervals estimated using Wilson’s continuity correction. P-value is from a one-tailed Fisher-exact test comparing SCIRA to DE. Also shown are the SE values for SCENIC, and SCENIC without the motif-enrichment analysis step (denoted “GENIE3”).

In contrast, if we were to only use the measured expression values of the TFs, we would have concluded that only 1 of the TFs (*Epas,* **Fig.3B**) exhibits upregulation. Of note, 4 of the 38 lung-specific TFs (*Gata2, Sox18, Tfec* and *Tnxb*) exhibited a 100% dropout rate across the 201 single cells, yet according to SCIRA all 4 exhibited some increase in activity during the developmental time course, in line with their known role in lung development. For instance, according to the mouse ENCODE transcriptomics dataset, adult lung is the tissue exhibiting highest expression of *Sox18* ^49^ (**SI fig.S8),** yet according to the FPKM data of Treutlein et al (**Fig.3B**) it is not expressed at all across the 201 single cells.

Given the overall discordance between the differential activity patterns derived from SCIRA and the differential expression values for the corresponding TFs, we decided to check that this was not due to our normalization procedure, or due to our DE analysis method. Thus, we selected 5 genes which, according to Treutlein et al ^17^, are significantly upregulated in mature (adult) alveolar type-2 cells compared to early progenitors (E14), and 5 which are significantly downregulated: we found that all 10 genes exhibited the expected differential expression pattern according to our normalized data and DE calling method (**SI fig.S9**). The discrepancy between the bulk RNA-Seq dataset on the one hand, and the scRNA-Seq data from Treutlein on the other, is therefore most likely due to the high technical dropout rate in the scRNA-Seq assay, which as we have seen may affect even highly expressing TFs such as *Sox18.* Thus, SCIRA is able to correctly infer that TFs like *Sox18* are highly active/expressed in the mature/adult alveolar cells, whilst direct inspection of the FPKM data does not (**Fig.3A**).

### Increased sensitivity of SCIRA over SCENIC

Next, we compared SCIRA to SCENIC ^30^, a recently proposed state-of-the-art algorithm for infering regulatory activity of transcription factors in single-cells. SCENIC uses the DREAM award-winning GENIE3 algorithm ^38^ to infer the regulatory network, and in contrast to SCIRA, it applies the network inference algorithm to the scRNA-Seq data itself. We reasoned that since SCENIC/GENIE3 relies on expression covariation across single cells to infer regulatory modules, that it would not be able to infer reliable regulons for many of the lung-specific TFs since many of these exhibit either high dropout rates or their expression values may be unreliable. Application of SCENIC to the Treutlein lung scRNA-Seq set resulted in motif-enriched regulatory modules for only 4 of the 38 lung-specific TFs (*Cebpd, Foxa2, Foxa1, Klf4*), yet these regulons failed to yield increased activity with time (**Methods**, **SI fig.S10**), resulting in zero sensitivity (**Fig.3E**). Because the motif-enrichment analysis was only significant for 4 of the 38 TFs, we next removed this step from SCENIC and defined regulons for each of the 38 TFs using only the first step in SCENIC, i.e. using GENIE3 to construct regulons based on covariation in expression only. However, this also resulted in zero sensitivity (**Fig.3E**). SCENIC results for liver were similar (**Fig.2E**). These results show that inferring regulons from scRNA-Seq data is not possible for many TFs that ought to exhibit clear dynamic activity patterns.

### SCIRA predicts inactivation of lung-specific TFs in lung tumor epithelial cells

To test SCIRA in a disease setting and to further show how it can reveal novel insight, we applied it to a recent lung cancer a scRNA-Seq study ^50^ which profiled a total of 52,698 cells (10X Chromium) derived from 5 lung cancer patients. Specifically, we hypothesized that many of our previously identified 38 lung-specific TFs would be inactivated in lung epithelial tumor cells ^37, 51^, since lack of differentiation is a well-known cancer hallmark ^52^. Following ^50^, we first categorized specific clusters of cells as alveolar (n=1709) and tumor epithelial (n=7450) (**Fig.4A**). We verified that the alveolar cells expressed relatively high levels of an alveolar marker (*CLDN18*) (**Fig.4B**), whilst both alveolar and tumor epithelial cells expressed relatively high levels of *EPCAM*, a well-known epithelial marker (**Fig.4C**). The great majority of alveolar cells were from non-malignant specimens ^50^, hence these cells largely represent normal (squamous) epithelium. Using SCIRA, we thus estimated regulatory activity for all 38 lung-specific TFs in each of the (1709+7450) cells, and computed t-statistics of differential activity between the alveolar and tumor epithelial cells. Remarkably, 35 out of the 38 TFs exhibited a statistically significant (Bonferroni adjusted P < 0.05) reduction in regulatory activity in the tumor cells (**Fig.4D**, **Wilcox-test P<1e-8**). Using 1000 Monte-Carlo randomizations, we verified that this number of inactivated TFs could not have arisen by chance (**Fig.4D, Monte Carlo P<0.001**). Among the most significantly inactivated TFs, we observed *FOXA2*, a TF required for alveolarization ^53, 54^ and that regulates airway epithelial cell differentiation (**Fig.4E**), and *NKX2-1*, a master TF of early lung development (**SI fig.S11**). Both of these TFs have been found to be inactivated in bulk tumor samples, representing lung carcinoma in situ (LCIS) ^37, 55^ and lung squamous cell carcinoma (LUSC) ^37^, consistent with SCIRA’s predictions. Other important TFs predicted by SCIRA to be inactivated included *SOX13,* which has been broadly implicated in lung morphogenesis ^56^, and *HIF3A* which has been shown to be highly expressed in alveolar epithelial cells and thought to be protective of hypoxia-induced damage ^57^ (**SI fig.S10**). Of note, the aryl hydrocarbon receptor (*AHR*), which is a regulator of mucosal barrier function and activation of which enhances CD4+ T-cell responses to viral infections ^58, 59^, was also strongly inactivated in the tumor epithelial cells (**SI fig.S10**). We stress that the inactivation of these TFs in lung cancer has been previously demonstrated by us in bulk RNA-Seq lung tissue data from The Cancer Genome Atlas (TCGA) ^37^, and therefore the prediction by SCIRA that they are also inactivated in single cells from the lung tumor epithelium not only provides further validation of SCIRA, but also illustrates its power to unravel novel biology: for instance, the inactivation of *HIF3A* and *AHR* in single cells from the tumor epithelium confirms that the inactivation observed in bulk tissue samples is indeed occurring in the epithelial compartment and not the result of confounding by stromal cells.

**Figure-4:**
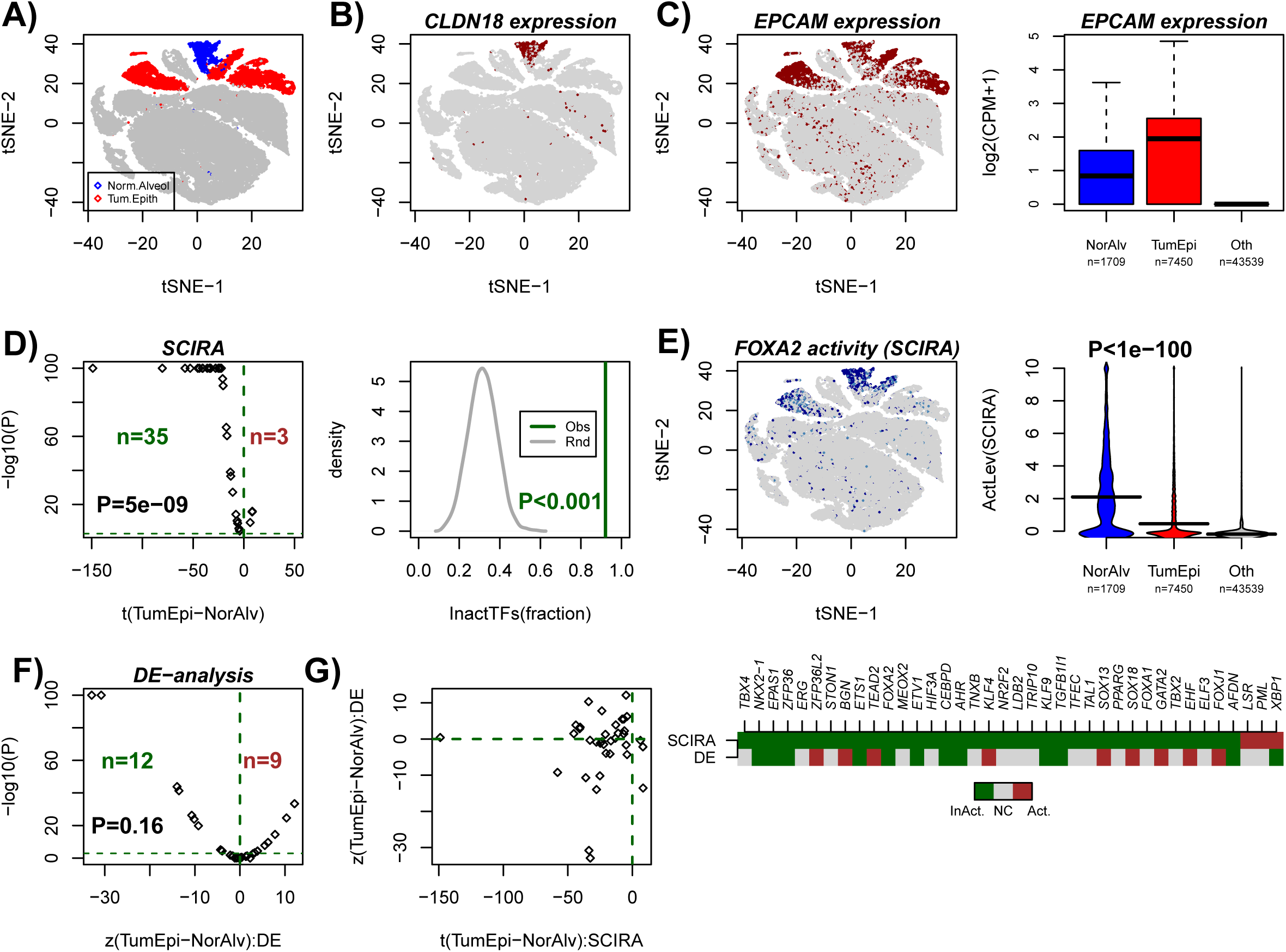
SCIRA predicts inactivation of lung-specific TFs in lung tumor epithelial cells. **A)** t-SNE scatterplot of approximately 52,000 single cells from 5 lung cancer patients, with a common non-malignant alveolar and (tumor) epithelial clusters highlighted in blue and red, respectively. **B)** Corresponding t-SNE scatterplot with cells colored-labeled by expression of an alveolar marker *CLDN18.* **C)** As B), but with cells colored according to expression of the epithelial marker *EPCAM.* Right panel depicts boxplots of the log_2_(counts per million + 1) of *EPCAM* for cells in the non-malignant alveolar cluster, the tumor epithelial clusters and all other cell clusters combined (T-cells, B-cells, endothelial, myeloid and fibroblast cells). **D)** Scatterplot of –log_10_(P-value) (y-axis) vs. the corresponding t-statistic (x-axis) of differential activity (as estimated using SCIRA) between tumor epithelial and normal alveolar cells for the 38 lung-specific TFs. P-values have been capped at P=1e-100. TFs passing a Bonferroni adjusted < 0.05 threshold and exhibiting decreased or increased activity in tumor epithelial cells indicated in darkgreen and darkred, respectively. P-value is from a one-tailed Wilcox rank sum test, testing whether the t-statistic distribution has a median significantly less than 0. Right panel depicts the Monte-Carlo (n=1000 runs) significance analysis with grey curve denoting the null distribution for the fraction of TFs exhibiting significant inactivation in tumor epithelial cells, and darkgreen line labeling the observed fraction (0.92=35/38). Empirical P-value derived from the 1000 Monte-Carlo runs is given. **E)** Scatterplot as in A), but now with cells color-labeled according to activation of *FOXA2* as estimated using SCIRA. Beanplots of the predicted SCIRA activity level of *FOXA2* between normal alveolar, tumor epithelial and all other cells. P-value is from a t-test between normal alveolar and tumor epithelial cell clusters. **F)** Scatterplot of –log_10_(P-value) (y-axis) vs. the corresponding z-statistic value (x-axis) of differential expression between tumor epithelial and normal alveolar cells for the 38 lung-specific TFs, where the P-value derives from a Wilcoxon rank sum test and where the z-statistic was derived by transforming the Wilcoxon rank sum test P-value into a quantile according to a standard normal distribution. P-values have been capped at P=1e-100. The number of TFs passing a Bonferroni adjusted < 0.05 threshold and exhibiting decreased or increased expression in tumor epithelial cells are indicated in darkgreen and darkred, respectively. P-value in plot is from a one-tailed Wilcox rank sum test, testing whether the z-statistic distribution has a median significantly less than 0. **G)** Scatterplot of the t-statistic of differential activity according to SCIRA (x-axis) vs. the z-statistic from DE analysis (y-axis) for all 38 TFs. Heatmap compares the predicted pattern of activation/inactivation of all 38 TFs according to SCIRA and DE analysis. Darkgreen denotes significant inactivation or underexpression in tumor epithelial cells compared to normal alveolar, brown denotes significant activation or expression. Grey=no-change (NC).

Importantly, most of the above insights would not have been obtained had we performed DE analysis on the 38 TFs (**Fig.4F-G, Methods**). For instance, according to a Wilcoxon rank sum test, 21 TFs were differentially expressed between the alveolar and tumor epithelial cells, but with no clear trend towards underexpression in tumor epithelial cells (**Fig.4F**). In addition, TFs such as *TBX4*, *SOX18* and *FOXJ1*, all with important roles in lung tissue development were not inactivated in tumor epithelial cells according to single-cell DE analysis, but were strongly inactivated according to SCIRA (**Fig.4G**). Given that all three TFs *TBX4*, *SOX18* and *FOXJ1* have been found to be inactivated/underexpressed in bulk samples representing LCIS and LUSC ^37^, this further confirms the improved sensitivity of SCIRA over ordinary DE analysis.

## Discussion

It has recently been argued that inferring regulatory effects from scRNA-Seq data is highly problematic due to their noisy nature and high dropout rates ^33^. The analysis presented here strongly supports this view. Indeed, by leveraging the power of the high-quality large scale bulk RNA-Seq dataset from the GTEX consortium, and by adjusting for cell-type heterogeneity to identify with high-confidence tissue-specific TFs, we have seen how many of these TFs were either subject to surprisingly high dropout rates, or did not exhibit the expected expression patterns in the corresponding differentiation time course scRNA-Seq datasets. We note that the overall dropout rates in the lung and liver scRNA-Seq sets considered here were 85% and 78%, respectively, and interestingly the original studies presenting these datasets did not remark how many of the known tissue-specific TFs (e.g. *Hnf1a* in liver, *Sox18* in lung) were not detected in their studies. Instead, these studies focused their discussion on other TFs which did exhibit the expected expression patterns. This therefore exposes a major limitation of methods that rely on DE analysis, including methods like SCENIC which aim to infer regulatory relations by correlating the expression patterns of the TFs to those of potential targets. Since the scRNA-Seq expression patterns of the TFs are unreliable, it is therefore in general not possible to infer reliable regulons, and correspondingly the observed dynamic TF-activity patterns as derived with SCENIC appeared to be largely random and were inconsistent with prior knowledge.

To overcome these limitations, we have here advocated an altogether different approach called SCIRA, demonstrating the feasibility of using predicted regulons of transcription factors as derived from cross-tissue bulk RNA-Seq data, to reliably infer transcription factor activity in single cells. Indeed, we have extensively validated this concept, and using SCIRA we were able to identify approximately 60% to 70% more TFs that play known key roles in differentiation, compared to SCENIC and standard DE analysis alone. Thus, the fact that the TF-regulons are derived from bulk data is not a severe limitation but a key advantage of our approach. This is in principle only true as long as the scRNA-Seq study in question profiles cells that are relatively abundant in the GTEX tissue set. However, we have seen that even in the case of liver-tissue where cholangiocytes make up a relatively small percentage (~5%) of all cells ^45^, SCIRA was still able to identify cholangiocyte specific TFs, in line with our power-calculation estimates. Indeed, SCIRA-derived TF activity patterns in single-cells recapitulated a bi-modal distribution with some TFs exhibiting a clear increase of activity in hepatocytes, while others exhibited the increase in cholangiocytes. Some of the latter TFs appear to be novel regulators of cholangiocyte fate (e.g. *Bgn, Irf6*), thus demonstrating that SCIRA can be used to identify novel dynamic patterns of regulatory activity at single-cell resolution, despite using regulons inferred from bulk data.

In addition, we have also shown how SCIRA can lead to critical novel insights in a disease context. The demonstration, at single cell resolution, that most of the lung-specific TFs are inactivated specifically in the lung tumor epithelial cells is entirely novel. Although highly consistent and expected given the analogous observations made at the bulk level (see e.g. ^37, 51^), it is not automatic that bulk and single-cell results would agree so well given the confounding effects of cell-type heterogeneity in bulk samples, specially in a tissue like lung where over 40% of cells are stromal cells ^60^. Many of the lung-specific TFs, which SCIRA predicts to be inactivated in lung tumor epithelial cells (e.g. *NKX2-1, FOXA2, FOXJ1*, *AHR, HIF3A*) implicate key cancer-pathways (lung differentiation, immune-response, hypoxia-response), and their inactivation likely represents key driver events. Supporting this, epigenetically induced silencing of *NKX2-1* has recently been proposed to be a key driver event in the development of lung cancer ^55^.

Finally, it is important to discuss in more detail the comparison between SCIRA and SCENIC. Based on our high-confidence tissue-specific regulatory networks, we estimated the sensitivity of SCENIC to detect true regulatory activity patterns to be less than 5%. However, SCENIC is an algorithm that attempts to infer regulons for all available TFs. In the case of human, assuming approximately 1500 TFs, a sensitivity of 5% means that we can identify the true dynamic activity patterns for approximately 75 TFs. Thus, application of SCENIC to a scRNA-Seq dataset can retrieve valuable patterns of regulatory activity, as shown by a number of studies ^30, 50^. However, as shown here, SCENIC can miss up to 95% of the most important TFs. On the other hand, we acknowledge that a main limitation of SCIRA is its tissue-centric approach, and therefore the need to build a tissue-specific regulatory network from adult tissue, which may therefore miss many key TFs that are expressed early-on in development, but which later, in adult tissue, are switched off. We acknowledge this, yet SCIRA could be easily extended to include additional bulk RNA-Seq datasets profiling samples from earlier time points in development and differentiation, to thus expand the repertoire of transcription factors. As such, SCIRA provides a flexible and easily generalizable framework in which to infer regulatory activity in single-cells.

In summary, our study has demonstrated that using TF-regulons can help overcome the high dropout rate of scRNA-Seq data, to enable accurate estimation of TF-activity in single cells. The quality of the TF-regulons is key to the reliability of the inference, and we have shown that high-quality approximate regulons can be constructed from large-scale bulk RNA-Seq datasets like GTEX. The computational algorithm SCIRA presented here, nicely complements methods like SCENIC, and is freely available as part of our SEPIRA Bioconductor package. SCIRA will be useful and broadly applicable to developmental time course scRNA-Seq studies, as well as those profiling single cells in diseases like cancer.

## Methods

### Single cell data and preprocessing

We analyzed scRNA-Seq data from a total of 3 studies:

*Treutlein Lung Differentiation set:* This scRNA-Seq Fluidigm C1 dataset derives from Treutlein et al ^17^. Normalized (FPKM) data were downloaded from GEO under accession number GSE52583 (file: GSE52583.Rda). Data was further transformed using a log2 transformation adding a pseudocount of 1, so that 0 FPKM values get mapped to 0 in the transformed basis. After quality control, there are a total of 201 single cells assayed at 4 different stages in the developing mouse lung epithelium, including embryonic day E14.5 (n = 45), E16.5 (n = 27), E18.5 (n = 83) and adulthood (n = 46).

*Yang Liver Differentiation set:* This scRNA-Seq Fluidigm C1 dataset was derived from Yang et al ^41^, a study of differentiation of mouse hepatoblasts into hepatocytes and cholangiocytes. Normalized (TPM) data were downloaded from GEO under accession number GSE90047 (file: GSE90047-Singlecell-RNA-seq-TPM.txt). Data was further transformed using a log2 transformation adding a pseudocount of 1, so that 0 TPM values get mapped to 0 in the transformed basis. After quality control, 447 single-cells remained, with 54 single cells at embryonic day 10.5 (E10.5), 70 at E11.5, 41 at E12.5, 65 at E13.5, 70 at 14.5, 77 at 15.5 and 70 at E17.5.

*Lung tissue and tumor microenvironment dataset:* This scRNA-Seq 10X Chromium dataset was derived from ^50^, a study profiling malignant and non-malignant lung samples from five patients. We downloaded all .Rds files available from ArrayExpress (E-MTAB-6149), which included the processed data and t-SNE coordinates, as well as cluster cell-type assignments. After quality control, a total of 52,698 single-cells remained of which 1709 were annotated as alveolar, 5603 as B-cells, 1592 as endothelial cells, 1465 as fibroblasts, 9756 as myeloid cells, 24911 as T-cells and 7450 as tumor epithelial cells. A small cluster of 212 cells was annotated as normal epithelial, yet they derived from a malignant sample ^50^, so given this inconsistency we removed these cells from any analysis, as according to us their “normal” nature is far from clear. The alveolar epithelial cell cluster derived mainly from non-malignant samples and was therefore considered most representative of the normal epithelial cells found in lung.

### GTEX bulk RNA-Seq dataset

We used the bulk RNA-Seq dataset from the GTEX resource ^36^. Specifically, the normalized RPKM data was downloaded from the GTEX website and annotated to Entrez gene IDs. Data was then log_2_ transformed with a pseudocount of +1. This resulted in a data matrix of 23929 genes and 8555 samples, encompassing 30 tissue types (adipose=577, adrenal gland=145, bladder=11, blood=511, blood vessel=689, brain=1259, breast=214, cervix uteri=11, colon=345, esophagus=686, fallopian tube=6, heart=412, kidney=32, liver=119, lung=320, muscle=430, nerve=304, ovary=97, pancreas=171, pituitary=103, prostate=106, salivary gland=57, skin=891, small intestine=88, spleen=104, stomach=193, testis=172, thyroid=323, uterus=83, vagina=96).

### The SCIRA algorithm

The SCIRA algorithm has two main steps: (i) construction of a tissue-specific regulatory network and (ii) inference of regulatory activity in single cells for the transcription factors (TFs) in the network constructed in step (i).

*(i) Construction of tissue-specific regulatory network:* For a given tissue-type, SCIRA infers a corresponding tissue-specific regulatory network using our SEPIRA algorithm ^37^. Briefly, SEPIRA implements a greedy partial correlation approach to infer a partial correlation network between regulators (TFs) and potential gene targets using the large GTEX cross-tissue dataset described above. The greedy partial correlation approach is similar in concept to the GENIE3 algorithm ^38^ (which was found to be one of the best performing reverse-engineering methods in the DREAM-5 challenge ^35^), in the sense that it infers the regulators for each gene in turn. SEPIRA however does this using partial correlations instead of trees. By computing partial correlations over 8555 samples across 30 different tissue-types, it is possible to identify direct regulatory relations that are relevant in the context of differentiation and development. Subsequently, SEPIRA identifies tissue-specific TFs, i.e. TFs with significantly higher expression in the given tissue type compared to all other tissues combined. This is done using the empirical Bayes moderated t-test framework (limma) ^61^. Importantly, a second limma analysis is performed by comparing the tissue of interest to individual tissue types if these contain cells that are believed to significantly infiltrate and contaminate the tissue of interest. Thus, in the case of liver we perform two limma analyses: comparing liver to all other tissue-types, and separately, liver to only blood, since blood contains immune cells which are known to infiltrate liver tissue accounting for approximately 40% of all cells found in liver ^60^. We require a liver-specific TF to be one with significantly higher expression in both comparisons. This ensures that the identified TFs are not driven by a higher immune cell (IC) infiltration in the tissue of interest compared to an “average” tissue where the IC infiltration may be low. As applied to liver, SEPIRA inferred a network of 22 liver-specific TFs and their regulons, with the average number of genes per regulon being 41, and with range 10 to 151. This network is available with this submission as an Rds file “netLIV.Rds” in **Supplementary File 1**.

The same procedure was used for lung, as described in our earlier work ^37^, resulting in a lung-specific network consisting of 38 lung-specific TFs and their regulons: the average number of genes in a regulon was 40, range was 10 to 152. This network is available with this submission as an Rds file “netLUNG.Rds” in **Supplementary File 2**.

*(ii) Estimation of regulatory activity:* Having inferred the tissue-specific TFs and their regulons, we next estimate regulatory activity of the TFs in each single cell of a scRNA-Seq dataset. This is done by regressing the log-normalized scRNA-Seq expression profile of the cell against the “target-profile” of the given TF, where in the target profile, any regulon member is assigned a +1 for activating interactions, a -1 for inhibitory interactions. All other genes not members of the TF’s regulon are assigned a value of 0. The TF-activity is then defined as the t-statistic of this linear regression. Before applying this procedure the normalized scRNA-Seq dataset is z-score normalized, i.e. each gene is centred and scaled to unit standard deviation.

We note that SCIRA relies on the high-quality tissue-specific regulatory network inferred in step-1. As such, SCIRA is particularly useful for scRNA-Seq studies that profile cells in the tissue of interest, either as part of a developmental or differentiation timecourse experiment, or in the context of diseases where altered differentiation is a key disease hallmark e.g. cancer and precursor cancer lesions.

### Power-calculation for SCIRA/SEPIRA

We used the following strategy to estimate the sensitivity of SEPIRA/SCIRA to detect highly expressed cell-type specific TFs in a given tissue, as a function of the corresponding cell-type proportion in the tissue. First, the main parameters affecting the power estimate include the relative sample sizes of the two groups being compared, the average expression effect size (in effect the average expression fold-change) of the cell-type specific TFs compared to all other cell-types, and the proportion of truly differentially expressed TFs. To estimate the sample sizes, we note that the median number of samples per tissue-type in GTEX is approximately 170. We therefore took a more conservative value of 150 to represent the number of samples in our tissue of interest, with the rest of samples in GTEX, i.e. 8555-150=8405, defining the number of samples from other tissue-types. For the proportion of truly differentially expressed TFs we made the conservative assumption that there are approximately 50 cell-type specific TFs, which by definition are therefore truly differentially expressed relative to all other cell-types from the other tissue-types. We note that while immune-cells are present across many different tissue-types, that SEPIRA already adjusts for this contamination ^37^. To estimate the average expression fold-change for top DEGs between single-cell types in a tissue, we analysed expression data from purified FACS sorted luminal and basal cells from the mammary epithelium ^62^. Because FACS sorted cell populations are still heterogeneous, we thus expect the resulting fold change estimates to be conservative. Using limma ^61^, we estimated the average expression fold-change to be 8 for the highest ranked DEGs, and approximately 6 among the top few hundred DEGs. Denoting the estimated fold change by FC, we then computed the effect size (*effS*) as *effS=log_2_[1*(1-mcf) + FC*mcf],* where *mcf* denotes the minor cell fraction, i.e. the proportion of cells in the tissue corresponding to the given cell-type. Finally, to estimate the sensitivity as a function of the significance level threshold, we used the parameter above as input to the TOC function of the OCplus R-package ^63^.

### Implementation of SCENIC

SCENIC is a pipeline of 3 distinct methods (GENIE3, RcisTarget, AUCell), each with its own Bioconductor package. We used the following versions: GENIE3_1.4.0, RcisTarget_1.2.0 and AUCell_1.4.1. Because the lung and liver scRNA-Seq sets are from mice, we used as regulators a list of 1686 mouse TF from the RIKEN lab (http://genome.gsc.riken.jp/TFdb/) together with the homologs of the human TFs in our lung and liver specific networks if these were not in the RIKEN lab list. GENIE3 was run with default parameter choices (treeMethod=”RF”, K=”sqrt”, nTrees=1000) but on a reduced data matrix where genes with a standard deviation less than 0.5 were removed. Regulons of TFs were obtained from GENIE3 using a threshold on the importance measure of 0.01, and only positively correlated targets were selected using a Spearman correlation coefficient threshold > 0. In SCENIC, the targets are then scanned for enriched binding motifs using RcisTarget. We used the 7species.mc9nr feather files for both 500bp upstream of the TSS and also for a 20kb window centered on the TSS. Any enriched motifs in both analyses were combined to arrive at a single list of enriched motifs and associated TFs. We then found the overlap with the annotated TFs from GENIE3, and only those that overlapped were considered valid TF regulons. For these we then estimated a regulatory activity score using an approach similar to the one implemented in AUCell, but one that is threshold independent, and therefore an improvement over the method used in AUCell. Specifically, the activity score was defined as the AUC of a Wilcoxon rank sum test, whereby in each single cell, genes are first ranked in decreasing order of expression, and the AUC-statistic is then derived by comparing the ranks of the regulon (all positively correlated) genes to the ranks of all other genes.

### Differential Expression (DE) analysis

In this work we compare SCIRA to ordinary DE analysis, as implemented using a Wilcoxon rank sum test. The use of a non-parametric test, which is distribution assumption free, works well for scRNA-Seq with high dropout rates. When comparing statistics of differential activity from SCIRA to those from DE analysis, we transform Wilcoxon rank sum test P-values into z-statistics using a quantile normal distribution, taking into account the magnitude of the AUC value from the Wilcoxon test (i.e. AUC values > 0.5 correspond to higher expression in one group compared to other, whereas AUC < 0.5 represents the opposite case).

### Code Availability

SCIRA functions are part of the SEPIRA Bioconductor R-package, freely available from http://bioconductor.org/packages/SEPIRA.

### Data Access

Data analyzed in this manuscript is already publicly available from the following GEO (www.ncbi.nlm.nih.gov/geo/) accession numbers: GSE52583, GSE90047, and from ArrayExpress (www.ebi.ac.uk/arrayexpress) accession numbers: E-MTAB-6149.

## Acknowledgements

This work was supported by NSFC (National Science Foundation of China) grants, grant numbers 31571359 and 31771464 and by a Royal Society Newton Advanced Fellowship (NAF award number: 164914). We would also like to thank Peter Kharchenko and John Marioni for useful discussions.

