## Supplementary Information for "Leveraging high-powered RNA-Seq datasets to improve inference of regulatory activity in single-cell RNA-Seq data"

### SUPPLEMENTARY FIGURES

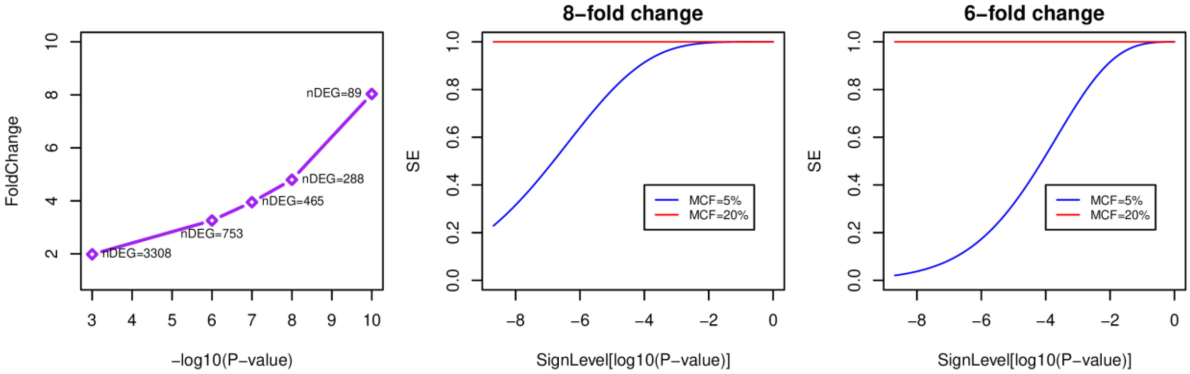


**fig.S1: Power-Analysis for SCIRA. Left panel:** Estimated Fold-Change (y-axis) between purified FACS sorted luminal and basal bulk samples from the mammary epithelium [1] against the significance-level (-$\log_{10}$[P-value], x-axis) with P-value derived from a moderated t-test. The number of differentially expressed genes (nDEG) at each significance threshold is given. Observe that 6 to 8-fold changes are not uncommon when comparing purified cell populations to each other. **Middle & Right panels**: Sensitivity (SE, y-axis) to detect putative transcription factors exhibiting 8 or 6-fold changes in expression in a cell-type with minor cell fraction (MCF) of 5% and 20% in the tissue (i.e. making up 5 and 20% of the cells in the tissue) against the significance level ($\log_{10}$[P-value], x-axis). The estimation is based using the GTEX dataset where the median number of samples per tissue-type is 150, and where the total number of samples, encompassing 30 tissue-types is 8555. Thus, the power analysis is performed for 150 samples vs 8405. In addition, we have assumed that there are 50 truly differentially expressed TFs within the tissue-type of interest.

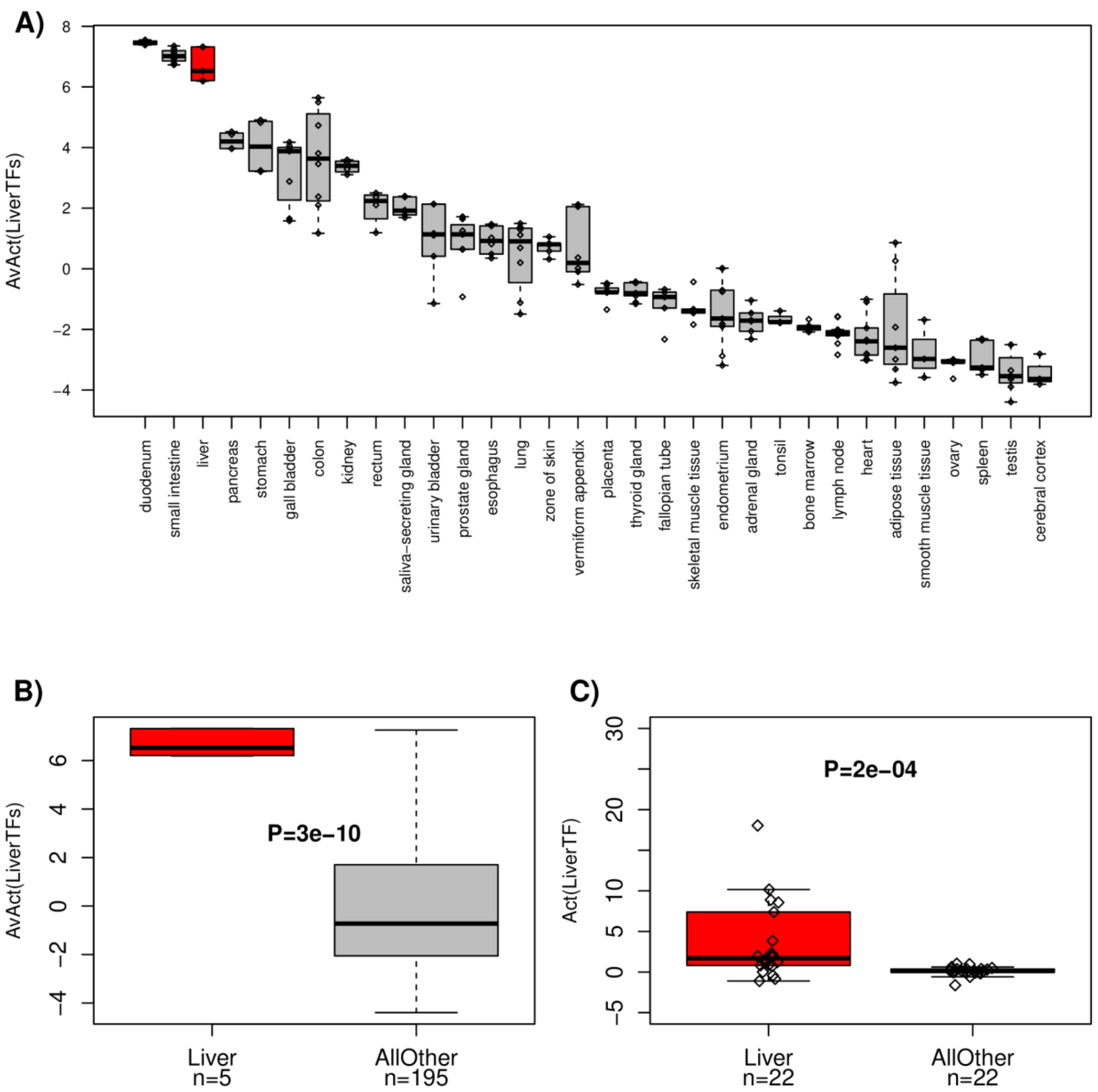


**fig.S2: Validation of liver-specific TFs and their targets in the multi-tissue RNA-Seq dataset from the Protein Atlas. A)** Boxplots of the average SEPIRA-estimated activity level of 22 liver-specific TFs across different tissues profiled as part of the Protein Atlas project. In red we highlight the tissue "liver". **B)** Comparison boxplot of the same average activity level between liver and all other tissues. Number of tissue samples in each group is given. P-value is from a one-tailed t-test. **C)** Boxplots of the individual 22 liver-specific TF activity levels in liver vs all other tissue-types, where values within a tissue have been averaged. P-value is from a one-tailed paired Wilcoxon rank sum test.

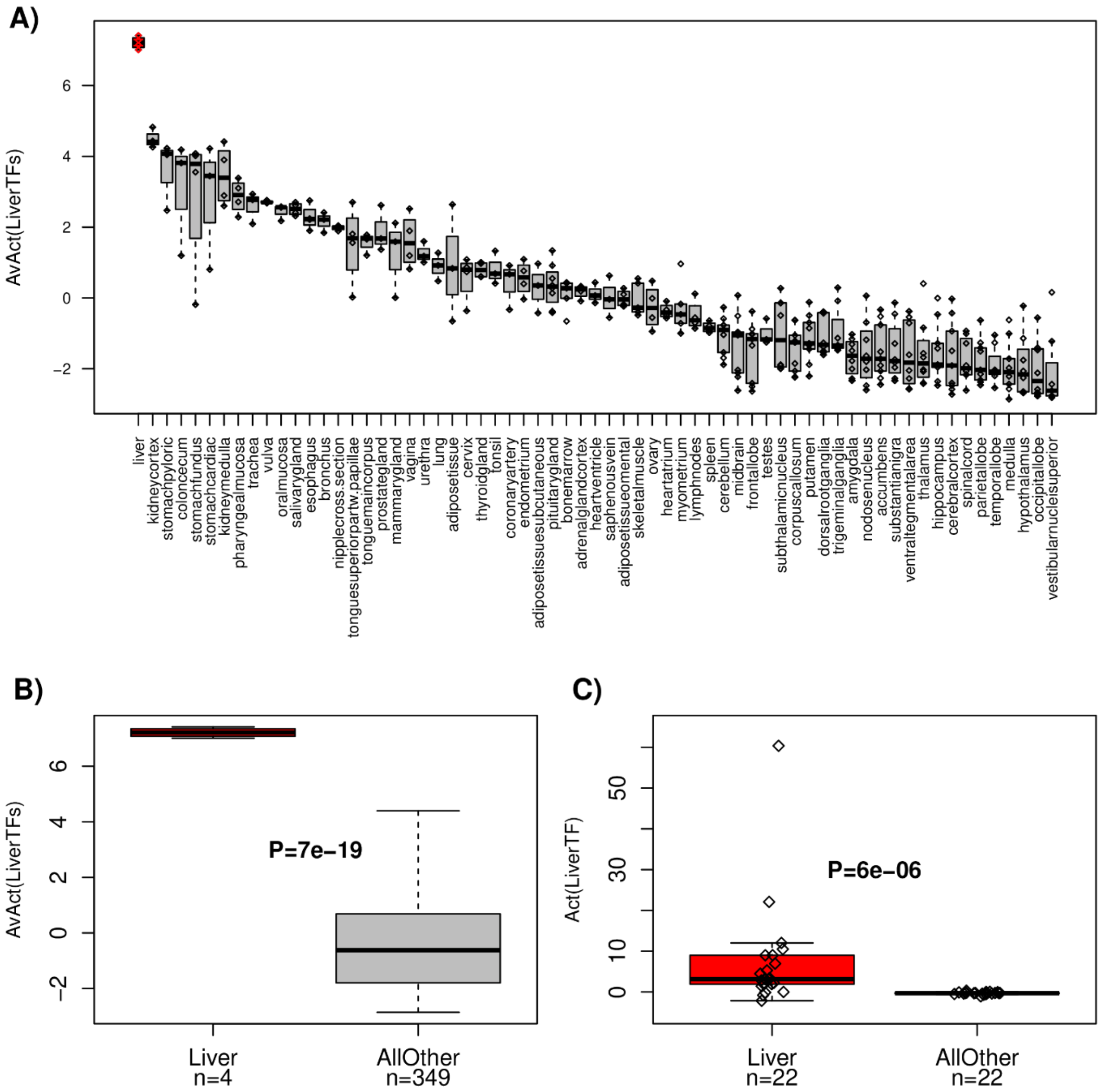


**fig.S3: Validation of liver-specific TFs and their targets in the Affymetrix multi-tissue mRNA expression data from Roth et al. A)** Boxplots of the average SEPIRA-estimated activity level of 22 liver-specific TFs across different tissues profiled in Roth et al [2]. In red we highlight the tissue "liver". **B)** Comparison boxplot of the same average activity level between liver and all other tissues. Number of tissue samples in each group is given. P-value is from a one-tailed t-test. **C)** Boxplots of the individual 22 liver-specific TF activity levels in liver vs all other tissue-types, where values within a tissue have been averaged. P-value is from a one-tailed paired Wilcoxon rank sum test.

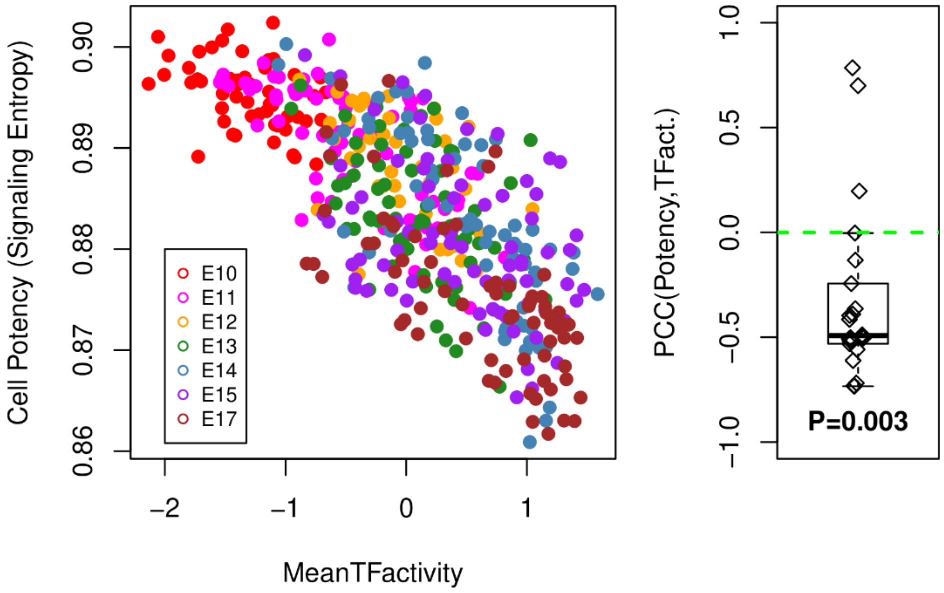


**fig.S4: Validation of SCIRA-derived TF activity estimates against an independent cell potency measure in the liver scRNA-Seq dataset.** Left panel depicts a scatterplot between the average TF-activity over the 22 liver-specific TFs (x-axis) and the cell potency measure as estimated using Signaling Entropy (y-axis) [3], with the 447 single cells from the Yang et al study [4] colored according to developmental timepoint. Right panel depicts the individual Pearson Correlation Coefficients (PCC) for each of the 22 liver specific TFs, where the PCC is computed between the TF-activity profile and the signaling entropy. Boxplot is shown and P-value is from a one-tailed Wilcoxon rank sum test.

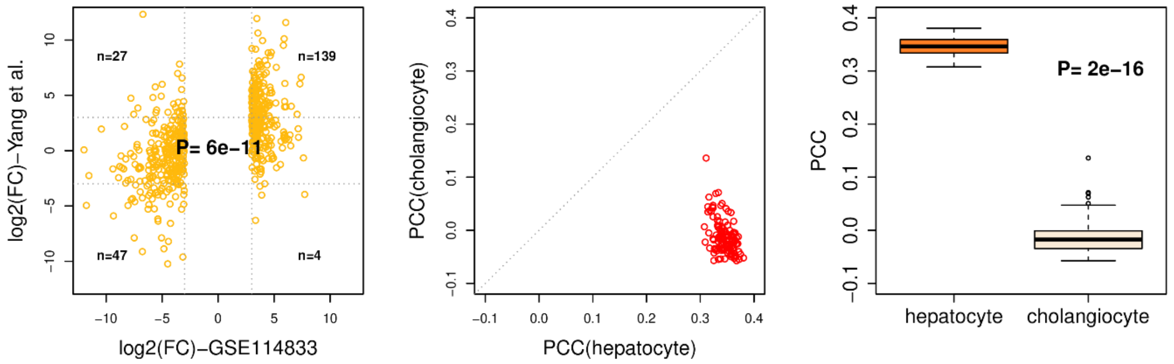


**fig.S5: Validation of hepatocyte/cholangiocyte expression signature and evaluation in GTEX bulk RNA-Seq dataset. Left panel:** Scatterplot of $\log_2$ fold-changes between hepatocyte and cholangiocyte samples from GSE114833 (x-axis) vs. their corresponding fold-changes in the scRNA-Seq data from Yang et al [4] for all genes called significant in the training set. Because only 1 hepatocyte and 1 cholangiocyte sample are available in GSE114833, only fold-changes could be used to determine significance. Number of data points in each significant quadrant are given. P-value is from a one-tailed Wilcox rank sum test. **Middle panel:** Scatterplot of the Pearson Correlation Coefficient (PCC) of the hepatocyte and cholangiocyte expression profiles constructed in GSE114883 using only differentially expressed genes with the corresponding bulk RNA-Seq expression profiles of the liver samples in GTEX. **Right panel:** Boxplot of the corresponding PCC values, demonstrating statistically significant higher PCC values for hepatocytes than cholangiocytes, and therefore that GTEX liver samples are composed mainly of hepatocytes. P-value is from a two-tailed Wilcox test.

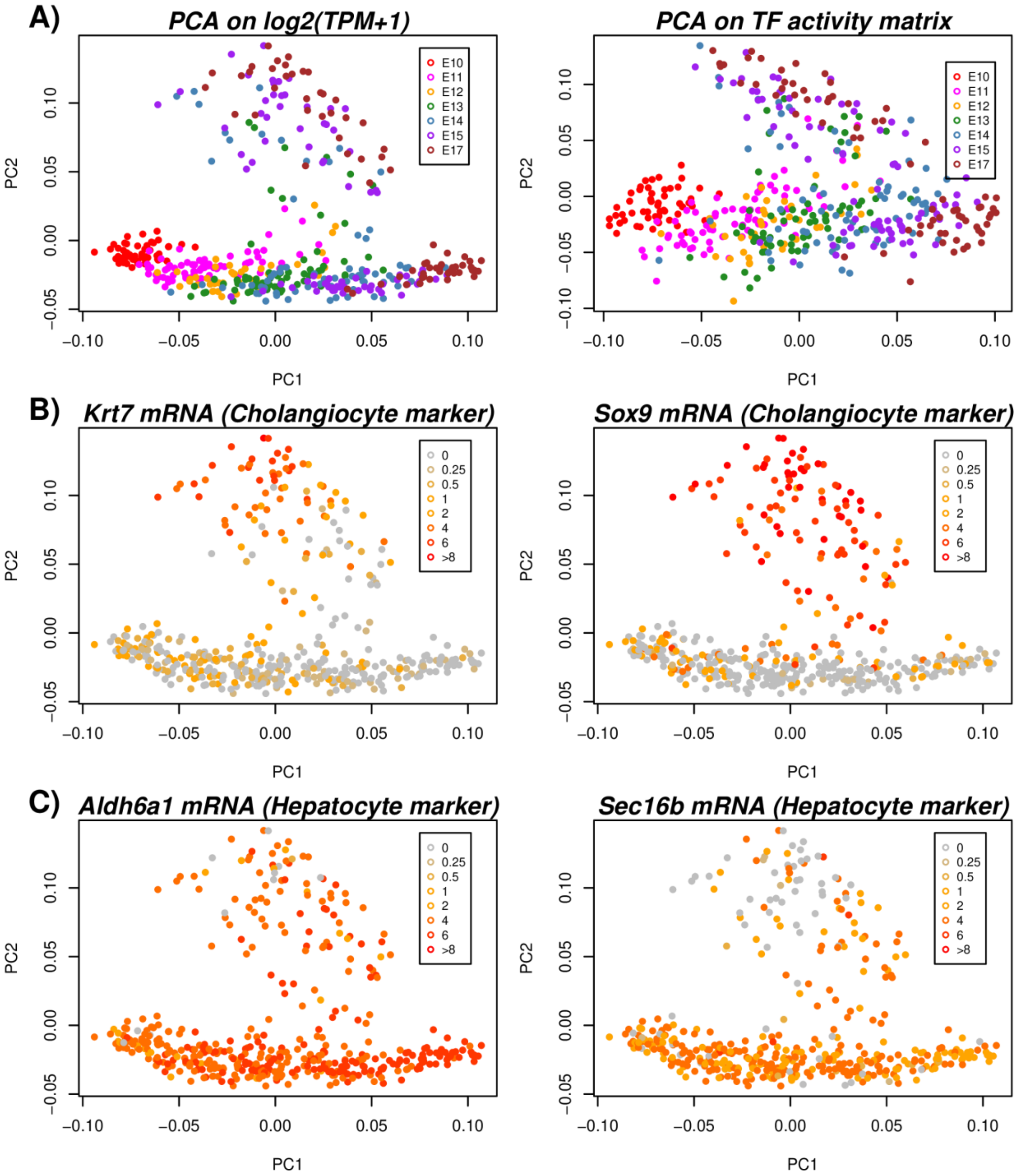


**fig.S6: PCA analysis on liver scRNA-Seq set and definition of cholangiocyte and hepatocyte branches. A)** Left panel depicts the PCA scatterplot of a PCA on the log2(TPM+1) expression matrix of the Yang et al liver scRNA-Seq study [4]. Right panel is the corresponding PCA scatterplot of a PCA on the transcription factor activity matrix over the 22 liver-specific TFs as estimated using SCIRA, which largely recapitulates the pattern derived from the full expression matrix. **B)** PCA scatterplot as in left panel of A), but now with cells colored according to the level of expression of two cholangiocyte markers (*Krt7*, *Sox9*) as identified in MacParland et al [5], defining the cholangiocyte branch. **C)** As B), but now for two hepatocyte markers (*Aldh6a1*, *Sec16b*), as identified in MacParland et al.

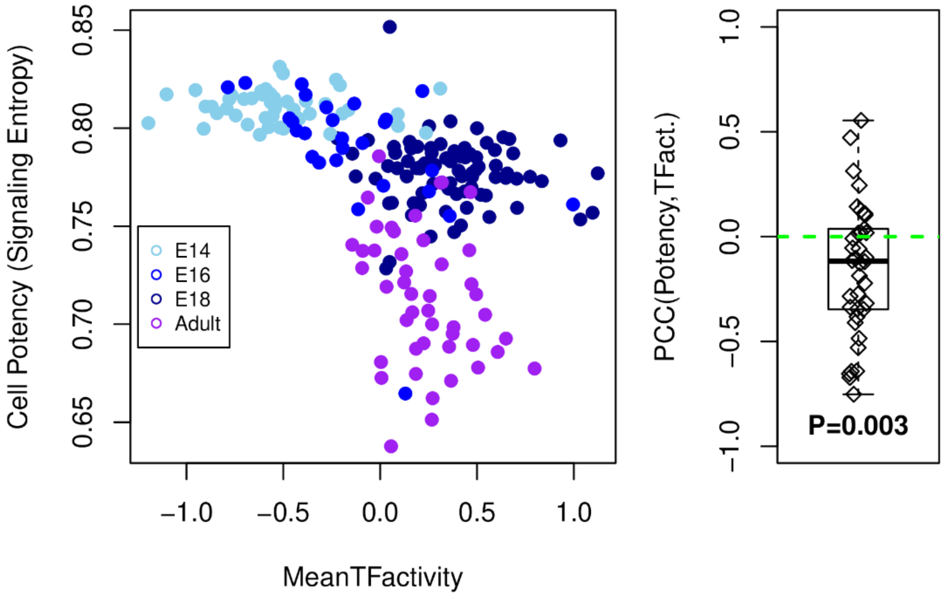


**fig.S7: Validation of SCIRA-derived TF activity estimates against an independent cell potency measure in the lung scRNA-Seq dataset.** Left panel depicts a scatterplot between the average TF-activity over the 38 lung-specific TFs (x-axis) and the cell potency measure as estimated using Signaling Entropy (y-axis) [3], with the 201 single cells from the Treutlein et al study [6] colored according to developmental time point. Right panel depicts the individual Pearson Correlation Coefficients (PCC) for each of the 38 lung specific TFs, where the PCC is computed between the TF-activity profile and the signaling entropy. Boxplot is shown and P-value is from a one-tailed Wilcoxon rank sum test.

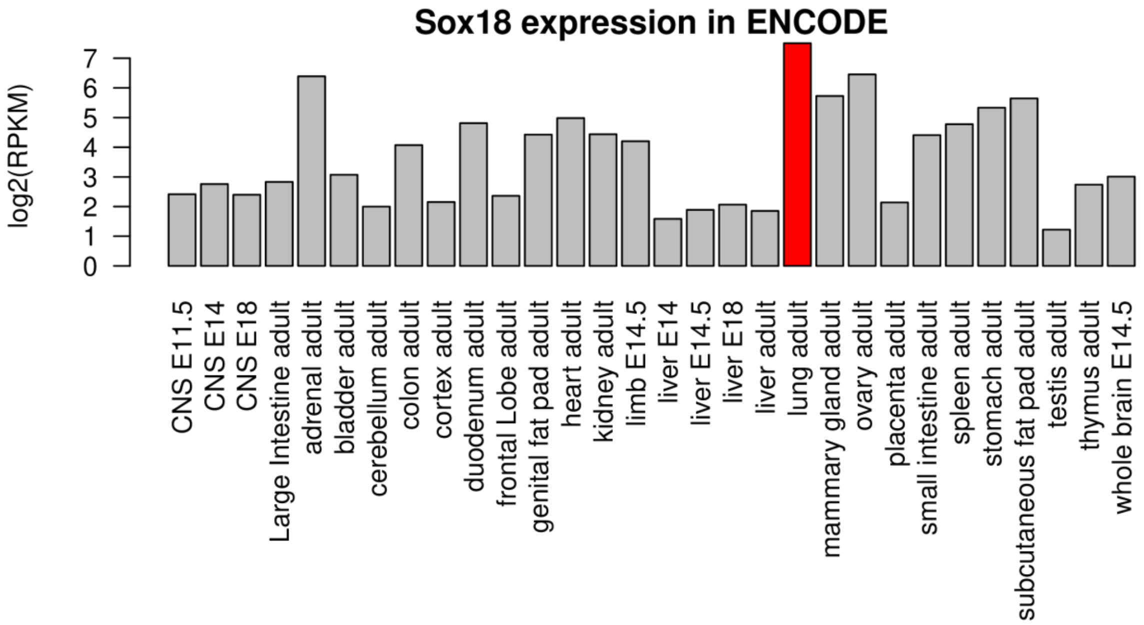


**fig.S8: Sox18 expression in the Mouse ENCODE transcriptomic dataset.** Barplot of *Sox18* expression ($\log_2$(RPKM)) in the mouse transcriptomic compendium of ENCODE, highlighting adult lung as the tissue of highest expression for *Sox18*.

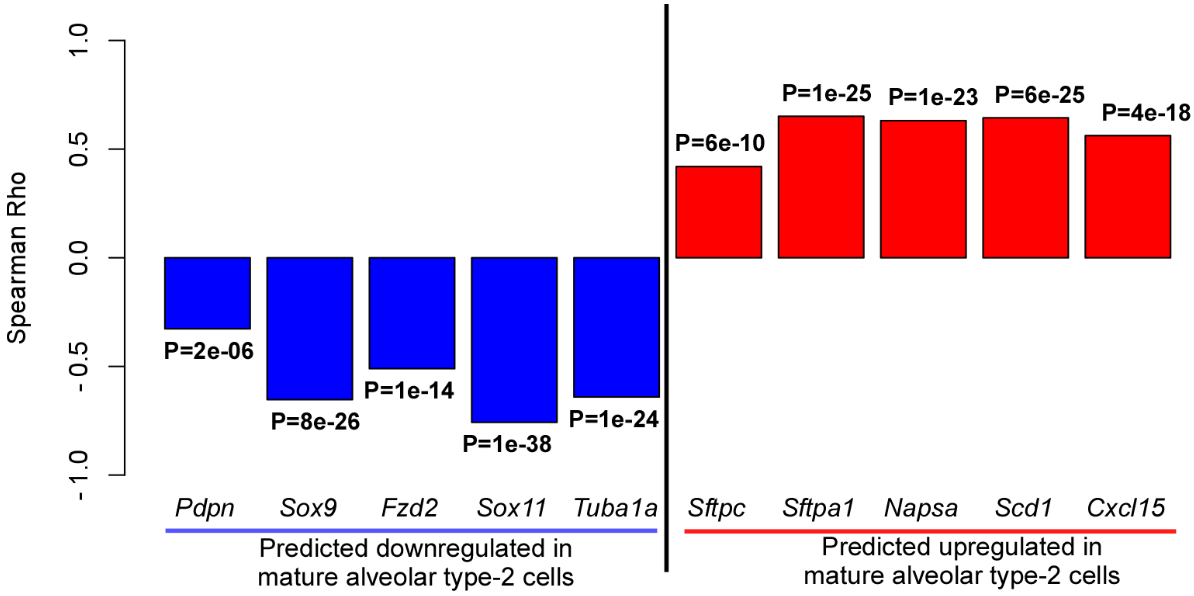


**fig.S9: Validation of DEG calling using Spearman Rank Correlation coefficient.** Barplots of Spearman rank correlation coefficients (rho) between log(FPKM+1.1) values and developmental time point (E14, E16, E18, Adult) in Treutlein et al data, for a total of 10 genes reported by Treutlein to exhibit differential expression between early progenitors (E14) and mature alveolar-type2 cells (Adult). According to Treutlein et al, the first 5 depicted genes exhibit downregulation in the adult cells, whereas the other 5 exhibit upregulation. The Spearman rank correlation analysis confirms this. P-values are from the Spearman rank-test.

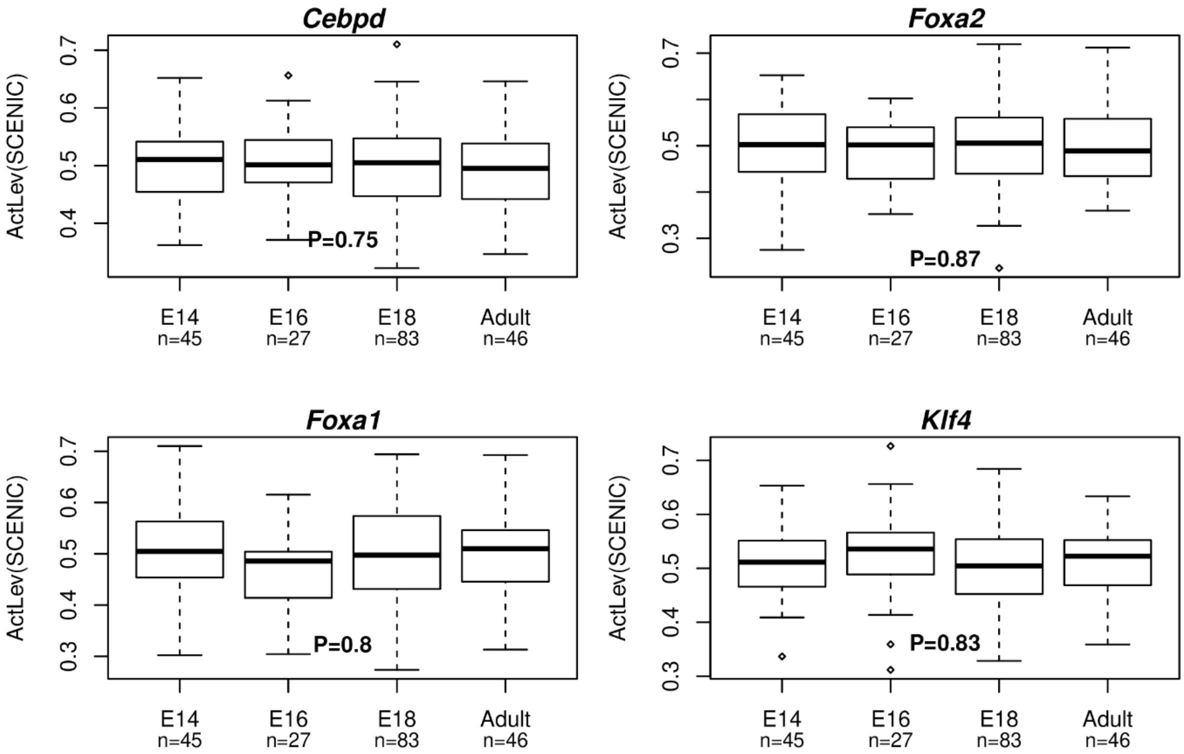


**fig.S10: SCENIC analysis in Treutlein lung scRNA-Seq dataset.** Boxplots of SCENIC inferred TF activity levels vs developmental timepoint in the Treutlein et al dataset for the 4 lung-specific TFs (*Cebpd*, *Foxa1*, *Foxa2*, *Klf4*) for which SCENIC inferred regulons that were enriched for corresponding TF binding motifs. P-value and t-statistic from a linear regression are given.

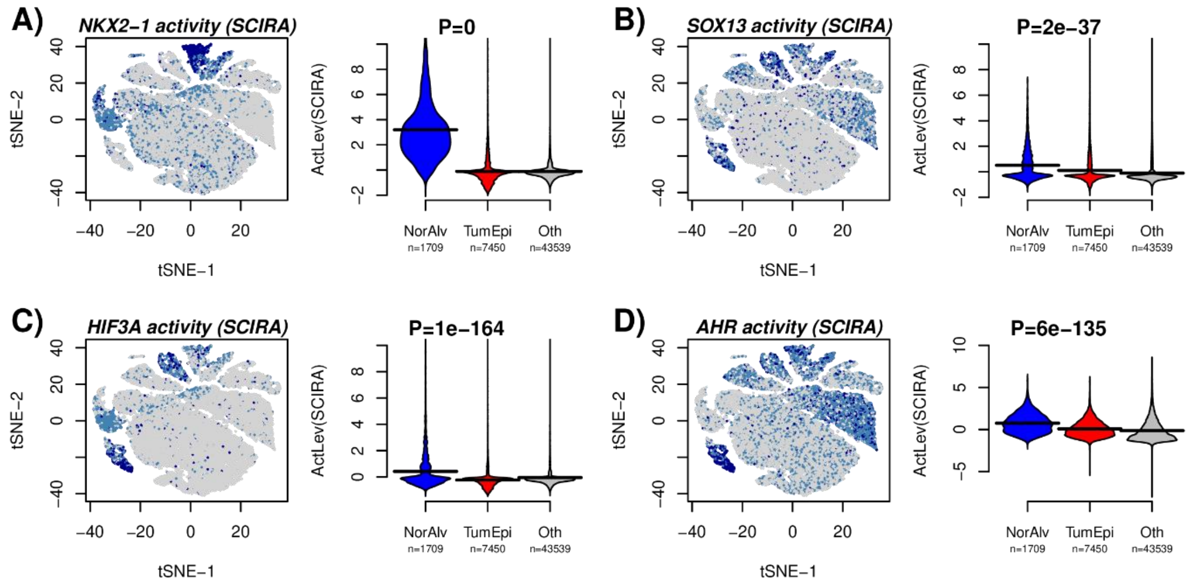


**fig.S11: Inactivation of lung-specific TFs in lung tumor epithelial cells. A)** t-SNE scatterplot of approximately 52,000 single cells from 5 lung cancer patients, with cells color-labeled according to the SCIRA predicted activity of *NKX2-1*. Right panel shows beanplots of the predicted SCIRA activity level of *NKX2-1* between normal alveolar, tumor epithelial and all other cells. P-value is from a linear model with activity level as response and normal alveolar/tumor epithelial status as predictor. P=0 means P<1e-500. **B-D)** As A), but now for the other lung-specific TFs *SOX13, HIF3A* and *AHR*.

### SUPPLEMENTARY TABLES

| TF(Human) | TF(Mouse) | # in Regulon | Positive | Inhibitory |
|---|---|---|---|---|
| *XBP1* | *Xbp1* | 18 | 18 | 0 |
| *LSR* | *Lsr* | 72 | 72 | 0 |
| *HNF4A* | *Hnf4a* | 31 | 28 | 3 |
| *BGN* | *Bgn* | 93 | 93 | 0 |
| *FOXA1* | *Foxa1* | 10 | 10 | 0 |
| *ONECUT1* | *Onecut1* | 15 | 15 | 0 |
| *HNF1A* | *Hnf1a* | 21 | 21 | 0 |
| *IRF6* | *Irf6* | 53 | 53 | 0 |
| *TIMELESS* | *Timeless* | 29 | 29 | 0 |
| *MYCL1* | *Mycl* | 40 | 38 | 2 |
| *TRIM15* | *Trim15* | 12 | 11 | 1 |
| *HNF4G* | *Hnf4g* | 37 | 28 | 9 |
| *FOXA2* | *Foxa2* | 15 | 15 | 0 |
| *NR1I2* | *Maats1* | 44 | 42 | 2 |
| *NR1I3* | *Nr1i3* | 151 | 151 | 0 |
| *ZKSCAN1* | *Zkscan1* | 23 | 23 | 0 |
| *TFB2M* | *Tfb2m* | 101 | 101 | 0 |
| *LHX2* | *Lhx2* | 13 | 12 | 1 |
| *ELF3* | *Elf3* | 71 | 71 | 0 |
| *BCL3* | *Bcl3* | 26 | 26 | 0 |
| *ZNF444* | *Zfp444* | 23 | 23 | 0 |
| *NR1H3* | *Nr1h3* | 10 | 10 | 0 |

**table.S1: Summary of the liver-specific regulatory network.** Table lists the human gene symbol and mouse homolog of the 22 liver-specific transcription factors (TFs) for the liver-specific regulatory network derived using the SEPIRA algorithm [7] on the GTEX dataset [8]. The other columns list the number of gene targets in the TF-regulon, and the number of these that represent positive and inhibitory interactions.

| TF(Human) | TF(Mouse) | # in Regulon | Positive | Inhibitory |
|---|---|---|---|---|
| *TFEC* | *Tfec* | 33 | 33 | 0 |
| *TBX2* | *Tbx2* | 18 | 14 | 4 |
| *FOXA2* | *Foxa2* | 15 | 15 | 0 |
| *TAL1* | *Tal1* | 19 | 19 | 0 |
| *TBX4* | *Tbx4* | 16 | 16 | 0 |
| *NKX2-1* | *Nkx2-1* | 24 | 24 | 0 |
| *GATA2* | *Gata2* | 13 | 13 | 0 |
| *EPAS1* | *Epas1* | 85 | 83 | 2 |
| *FOXJ1* | *Foxj1* | 152 | 152 | 0 |
| *LDB2* | *Ldb2* | 63 | 63 | 0 |
| *ETS1* | *Ets1* | 35 | 35 | 0 |
| *ETV1* | *Etv1* | 11 | 11 | 0 |
| *ERG* | *Erg* | 44 | 44 | 0 |
| *ELF3* | *Elf3* | 71 | 71 | 0 |
| *SOX13* | *Sox13* | 14 | 14 | 0 |
| *AHR* | *Ahr* | 39 | 38 | 1 |
| *PML* | *Pml* | 33 | 28 | 5 |
| *FOXA1* | *Foxa1* | 10 | 10 | 0 |
| *MLLT4* | *Mllt4* | 26 | 26 | 0 |
| *BGN* | *Bgn* | 93 | 93 | 0 |
| *ZFP36* | *Zfp36* | 19 | 18 | 1 |
| *TNXB* | *Tnxb* | 40 | 40 | 0 |
| *SOX18* | *Sox18* | 60 | 60 | 0 |
| *TEAD2* | *Tead2* | 53 | 52 | 1 |
| *XBP1* | *Xbp1* | 18 | 18 | 0 |
| *MEOX2* | *Meox2* | 42 | 41 | 1 |
| *KLF4* | *Klf4* | 20 | 20 | 0 |
| *HIF3A* | *Hif3a* | 10 | 10 | 0 |
| *LSR* | *Lsr* | 70 | 70 | 0 |
| *KLF9* | *Klf9* | 15 | 15 | 0 |
| *STON1* | *Ston1* | 31 | 30 | 1 |
| *PPARG* | *Pparg* | 16 | 16 | 0 |
| *ZFP36L2* | *Zfp36l2* | 24 | 24 | 0 |
| *CEBPD* | *Cebpd* | 17 | 17 | 0 |
| *TRIP10* | *Trip10* | 42 | 23 | 19 |
| *NR2F2* | *Nr2f2* | 31 | 24 | 7 |
| *TGFB1I1* | *Tgfb1i1* | 112 | 108 | 4 |
| *EHF* | *Elf3* | 77 | 50 | 27 |

**table.S2: Summary of the lung-specific regulatory network.** Table lists the human gene symbol and mouse homolog of the 38 lung-specific transcription factors (TFs) for the lung-specific regulatory network derived using the SEPIRA algorithm [7] on the GTEX dataset [8]. The other columns list the number of gene targets in the TF-regulon, and the number of these that represent positive and inhibitory interactions.
